# Dissociable mechanistic contributions of limb and task related errors during human sensorimotor learning

**DOI:** 10.1101/2020.11.13.381285

**Authors:** Anushka Oza, Adarsh Kumar, Apoorva Sharma, Pratik K. Mutha

## Abstract

The unpredictable nature of our world can introduce a variety of errors in our actions, including sensory prediction errors (SPEs) and task performance errors (TPEs). SPEs arise when our existing internal models of limb-environment properties and interactions become miscalibrated due to changes in the environment, while TPEs occur when environmental perturbations hinder achievement of task goals. The precise mechanisms employed by the sensorimotor system to learn from such limb- and task-related errors and improve future performance are not comprehensively understood. To gain insight into these mechanisms, we performed a series of learning experiments wherein the location and size of a reach target were varied, the visual feedback of the motion was clamped along fixed directions, and instructions were carefully manipulated. Our findings indicate that the mechanisms employed to compensate SPEs and TPEs are dissociable. Specifically, our results fail to support theories that suggest that TPEs trigger implicit refinement of reach plans, or that their occurrence automatically modulates SPE-mediated learning. Rather, TPEs drive improved action selection, that is, the selection of verbally-sensitive, volitional strategies that reduce future errors. Moreover, we find that exposure to SPEs is necessary and sufficient to trigger implicit recalibration. When SPE-mediated implicit learning and TPE-driven improved action selection combine, performance gains are larger. However, when actions are always successful and strategies are not employed, refinement in behaviour is smaller. Flexibly weighting strategic action selection and implicit recalibration could thus be a way of controlling how much, and how quickly, we learn from errors.

## Introduction

Humans often have to perform actions under challenging and changing conditions. For example, a golfer may have to tee off against a constant breeze, a dancer may be required to perform while wearing a new, heavier costume, or a patient may have to adjust their gait to compensate for an emerging neurological disorder. Understanding how we adapt our actions to such changes, and delineating the neural systems that support such learning has been a major goal in cognitive neuroscience. Laboratory tasks often simulate such perturbing conditions using various novel visual (1–3) or dynamic (4, 5) environments that induce errors in our movements. For instance, these perturbations can create a mismatch between the intended and actual sensory consequences of action, or sensory prediction errors (SPEs). SPEs are essentially limb-related execution errors that result from a miscalibrated internal model of the properties of the body and the environment. At the same time, perturbing environments can bring about a failure to accomplish the intended task goal (task performance errors, TPEs). While years of work has demonstrated we can learn to adjust our motor plans to account for such perturbation-induced errors, a gap exists in our understanding of the relative influence of various errors and the mechanisms they stimulate to enable such learning.

Experimental investigations and theoretical models suggest that SPEs drive iterative changes in motor plans by implicitly updating our internal models of the relationship between actions and their sensory consequences (6–8). This is not very different from ideas in predictive coding, which emphasizes that the brain generates and updates internal models of the world in order to improve future expectations (9). Implicit learning driven by SPEs has many interesting features: it evolves slowly (10), asymptotes similarly for different error magnitudes (11, 12), can be quite inflexible (13) and is impervious to verbal instruction (3, 6). There is also extensive evidence that learning from SPEs requires intact cerebellar and posterior parietal circuits. Cerebellar and parietal damage, or its disruption using brain stimulation techniques, impairs adaptation to visual (1–3, 14) and dynamic (15, 16) perturbations.

In contrast, controversy exists about how outcomes such as task success or failure influence the updating of action plans. One possibility is that performance failures, or TPEs, trigger intentional cognitive strategies to reduce perturbation-induced errors (17). For example, failure to obtain reward upon missing a target may make people consciously aware of the perturbation and so they might deliberatively adjust their aim to compensate for that error. However, alternative views have emerged from work examining the influence of binary reward on error-based learning (18–20). These studies have generally shown that learning is greater when TPEs occur compared to when they do not. Such findings can be explained by including a second, TPE-driven implicit process in computational models of learning (20, 21). Thus, in this framework, TPEs independently induce implicit learning, and this process additively combines with the SPE-mediated implicit component to determine the net change in action plans. A third possibility is that learning is actually only SPE-driven, but is modulated by TPEs (20, 22). Specifically, when a movement is unsuccessful, and hence unrewarded, a gain factor amplifies learning driven by SPEs (or alternatively, a positive reinforcement signal associated with a successful movement dials it down). Given these vastly varying hypotheses – one positing deployment of volitional strategies, another postulating the stimulation of an independent implicit process, and a third advocating for modulation – a clear understanding of the mechanistic effects of task outcomes has remained elusive.

In this work, we set out to address this gap and decisively probe how different error sources influence our planning of future actions. Our results fail to support the view that task-related performance failures, or TPEs, independently induce implicit learning; we rather note that implicit learning is driven only by limb-related SPEs. We also do not find evidence in favor of the view that TPEs modulate SPE-driven learning. Rather, our results advocate that TPEs trigger time-consuming strategic processes that are responsive to verbal instruction, and suggest that sustained task success reduces or eliminates strategy use during learning. Flexibly combining SPE-based implicit processes and TPE-driven strategic action selection could then be a way for the sensorimotor system to optimize how much and how rapidly we learn from errors.

## Results

### Experiment 1

In our first experiment, two groups of participants performed targeted reaching movements in three blocks: baseline, learning and washout. The hand was not directly visible during the reach, but visual feedback was provided by means of an on-screen cursor (Fig S1a). During each learning trial, the target was shifted or “jumped” to a location 10° counterclockwise to the original location. For one group (“Hit”), this jump was accompanied by an increase in target size; for the other group (“Miss”), the target size was not changed (Fig S1b). Critically, on the learning trials, motion of the feedback cursor was always clamped in the direction of the *original* target. This clamp ensured that the cursor always missed the new (jumped) target for the Miss group. This failure to strike the target with the cursor resulted in a TPE in the Miss participants. However, for the Hit group, the increase in target size ensured that the cursor always hit it, and thus, no TPE occurred. Importantly however, both groups experienced an SPE early on, which occurred as subjects directed the hand to the (center of) the new target (and expected the cursor to follow), but the cursor remained clamped in the direction of the original target. In sum, the Miss group experienced both an SPE and a TPE, while the Hit group experienced only an SPE.

When exposed to target shifts on learning trials, Hit participants began gradually aiming towards the new target but did not go outside the target envelope (Fig 1a). In contrast, the Miss participants (Fig 1b) quickly began directing the hand to the new target, but remarkably, continued to aim well beyond it, with some saturation emerging towards the end of the learning phase. This pattern was also seen in the group data (Fig 1c). During early learning, the mean hand deviation of the Miss participants was already greater than zero (mean = 3.870±1.727, CI = [0.166, 7.573]). Additionally, this deviation was greater than that seen in the Hit participants, but the group difference was not statistically reliable (Fig 1d; Hit mean: 0.910±0.630°, t(17.66) = −1.61, p = 0.125, Cohen’s *d* = −0.588) perhaps due to higher variability in the Miss participants’ reaches. The Miss group also showed a larger change in reaction time (RT) from late baseline levels compared to the Hit group (Fig 1e; t(18.94) = −3.07, p = 0.0063, Cohen’s *d* = −1.123). In fact, RT for Hit participants did not differ from baseline (t(14) = 0.25, p = 0.806, Cohen’s *d* = - 0.065).

**Fig 1.**
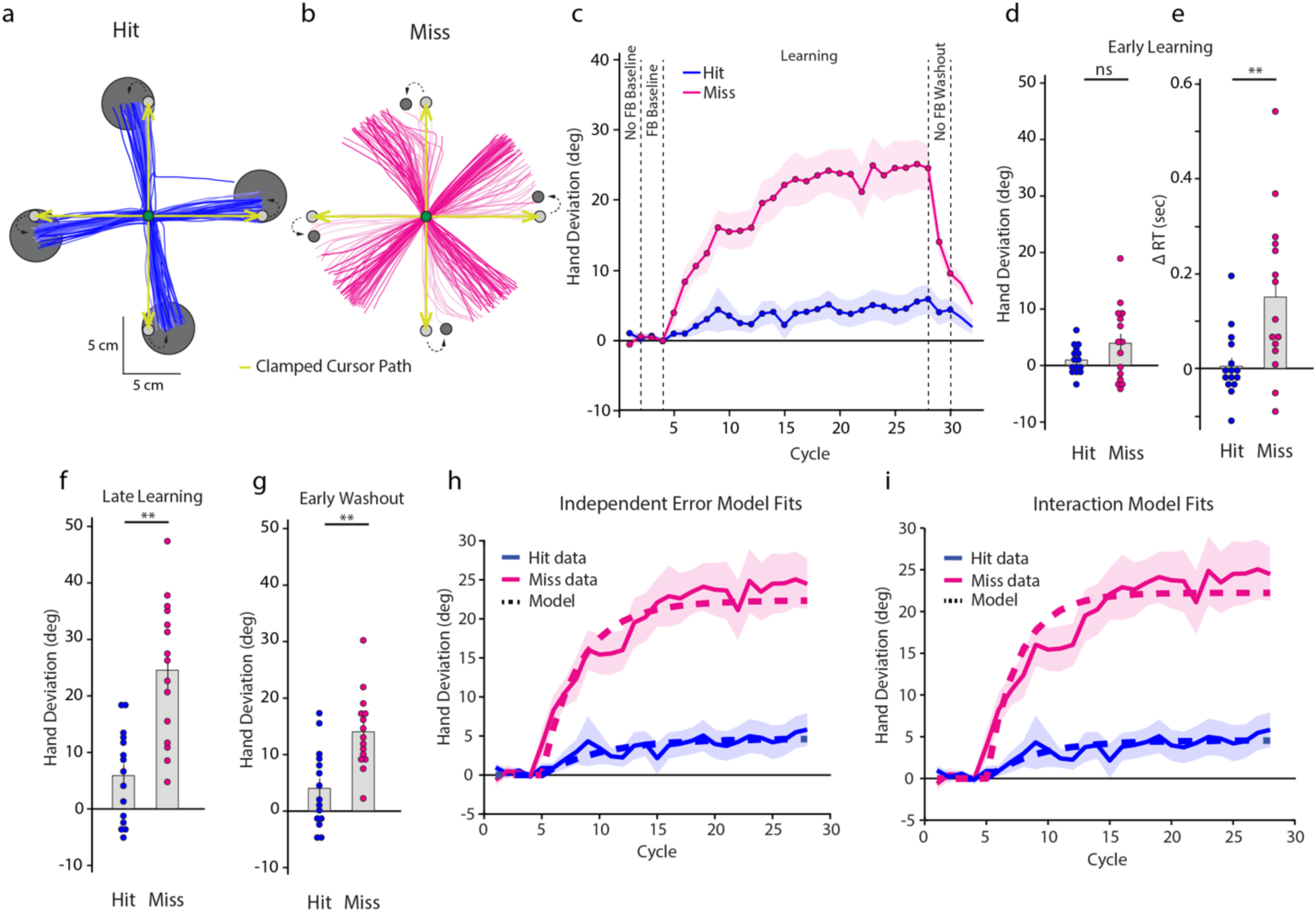
Hand trajectories of example subjects of the **(a)** Hit and **(b)** Miss groups during the learning block. Earlier trials are in lighter shades. The first 10 cm of the movement (corresponding to the target distance) are shown. The cursor path is indicated by green arrows. **(c)** Group averaged hand deviation (relative to the original target direction) across cycles of the experimental blocks. Shaded regions denote SEM. **(d)** Hand deviation and **(e)** change in RT during the early learning phase. **(f)** Hand deviation during the late learning and **(g)** early washout phases. Dots represent individual participants. **(h)** Independent error model and **(i)** Interaction model fits to the data of experiment 1. Solid lines represent the experimentally observed hand deviation while dashed lines represent the model fits. Shaded region denotes SEM.

As learning progressed, changes in reach direction occurred in both groups. However, by the end of learning, the Miss group had deviated much more (Fig 1f, t(23.99) = −4.878, p < 0.001, Cohen’s *d* = −1.781). Critically, it was not the case that the Hit group showed no change; a direct comparison between the early and late learning phases of this group revealed a clear increase in hand deviation (t(14) = −2.726, p = 0.016, Cohen’s *d*= −0.704). However, relative to Miss participants, this learning was conspicuously attenuated. Interestingly though, we found no relationship between the change in hand angle from late baseline to early learning, and the change over the remainder of the learning block (Fig S2a; Hit: R^2^ = 0.004, p = 0.812; Miss: R^2^ = 0.082, p = 0.3). The absence of such an association suggested that changes in hand angle beyond the initial stage were largely independent of changes that might have occurred early on.

During early washout (without cursor feedback), hand deviation continued to be large for the Miss group (13.978±1.740°) indicating the presence of after-effects. After-effects were also evident in the Hit group (3.960±1.792°, CI = [0.117, 7.803]) although they were smaller (Fig 1g; t(27.98) = −4.011, p < 0.001, Cohen’s d = −1.465). Importantly, these after-effects were sustained for an extended period, with both groups continuing to show large deviations even after cursor feedback was restored (Miss: 8.026±0.966°, CI = [5.955, 10.097]; Hit: 3.191±1.442°, CI = [0.1, 6.283]). In fact, for the Miss group, after-effects did not return to zero even after all washout trials had ended (mean = 5.114±1.036°, CI = [2.891, 7.336]).

### Mathematical Modeling

What drives the greater change in hand angle in the Miss compared (the group that experienced TPEs) to the Hit group (the group that doesn**’**t experience TPEs)? To address this, we considered two mathematical models that probe the influence of task-related errors on learning (20).

In the first “Independent Error” model, the presence of TPEs sets off an independent implicit learning process that combines with SPE-mediated implicit learning to produce the net change in motor output on a trial-by-trial basis. The equations governing the trialwise updates in this framework are:

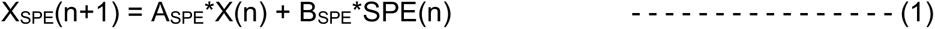

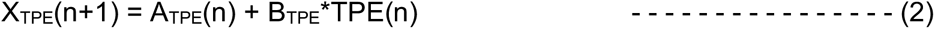

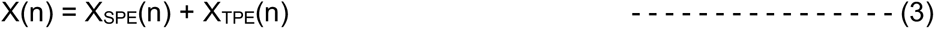

where X_SPE_(n) and X_TPE_(n) denote the state of SPE-driven and TPE-driven implicit learning components on the n^th^ trial respectively, while X(n) denotes the net motor output. A_SPE_ and A_TPE_ are retention factors that determine how much of the prior learning is carried over to the next trial, while B_SPE_ and B_TPE_ are learning rates for the SPE- and TPE-driven implicit mechanisms respectively.

In the second “Interaction” model, TPEs cannot by themselves induce implicit learning, but can only modulate implicit learning induced by SPEs. The equations governing the trial-by-trial updates to motor output then are:

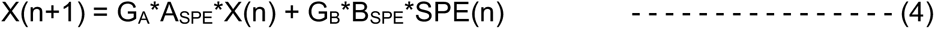

Here the state update is driven only by SPEs, but the modulation factors G_A_ and G_B_ (for A_SPE_ and B_SPE_ respectively) modulate this learning when TPEs are present. For fitting, the values of G_A_ and G_B_ are assumed to be one when TPEs are present, while they are allowed to vary when TPEs are absent.

#### Independent error model fit

We first fit the Independent Error model to the data from Experiment 1 (Fig 1h). To begin, since the Hit participants did not experience TPEs, only the SPE-based component (eq. 1) was fit to data from the Hit participants. We obtained a good fit (R^2^ = 0.81, RMSE = 0.7553) that yielded parameter estimates as A_SPE_ = 0.8184 and B_SPE_*SPE = 0.8441. We then used these estimates in the combined model that was fit to the data of the Miss group, and estimated the value of A_TPE_ and B_TPE_*TPE. Again, we obtained a good fit (R^2^ = 0.90, RMSE = 2.6833) with the parameter values as A_TPE_ = 0.7987, B_TPE_*TPE = 3.9622.

#### Interaction model fit

The interaction model assumes that the learning is only SPE driven, but the presence of a TPE brings about its modulation. To estimate model parameters, we fit this model to the data of the Miss group, while keeping G_A_ and G_B_ fixed at 1 (Fig 1i). We again obtained a good fit (R^2^ = 0.89, RMSE = 2.819, and parameter values were estimated as A_SPE_ = 0.6735, and B_SPE_*SPE = 7.2607. We then used these estimated parameter values to fit the model to the mean data of the Hit group and probe how G_A_ and G_B_ would change. We obtained R^2^ = 0.7938 and RMSE = 0.7724, while G_A_ and G_B_ were estimated to be 1.1534 (CI = [0.2147,1.4347]) and 0.1401 (CI = [0.0249,0.4913]) respectively (confidence intervals derived from fits to 10000 bootstrap samples of data from this group). Importantly, the confidence intervals of the parameter values indicated that while G_A_ was not different from 1, G_B_ differed significantly from this value. This indicated that substantial modulation of the SPE driven learning (the B_SPE_*SPE parameter) could occur via TPEs.

Thus, our modeling effort revealed that both, the Independent Error model as well as the Interaction model could account for the differences between the Hit and Miss groups of Experiment 1.

### Experiment 2

The success of both models in accounting for the results of Experiment 1 presented a conundrum. In particular, the success of the Independent Error model suggested that it was at least *mathematically* possible to account for the higher learning in the Miss group via an implicit mechanism triggered by TPEs that acts in concert with a different, SPE-driven implicit process. A pivotal *empirical* prediction of this framework is that since implicit learning is impervious to verbal instruction (3, 6), such learning should occur even when participants are explicitly instructed to ignore the shift in target location (TPE). We tested this hypothesis in our second experiment. We recruited two groups of participants that underwent training like the Miss group of Experiment 1 (TPEs present), but were told to ignore the (10° or 20°) jump and reach to the original target location.

What do our two models predict in these conditions? The Independent Error model suggests that learning would still progress because the TPE-based implicit process will not be suppressed by verbal instruction. In contrast, the Interaction model predicts no change in hand direction (Fig 2a). This is because in this framework, what drives learning is an SPE. In this experiment, since subjects are told to move in the direction of the original target, and the cursor also follows in the same direction, an SPE is not induced. That is, since there is no mismatch between the expected actual curosr motion, an SPE is absent. Under such conditions, the Interaction framework predicts no change in hand angle.

**Fig 2.**
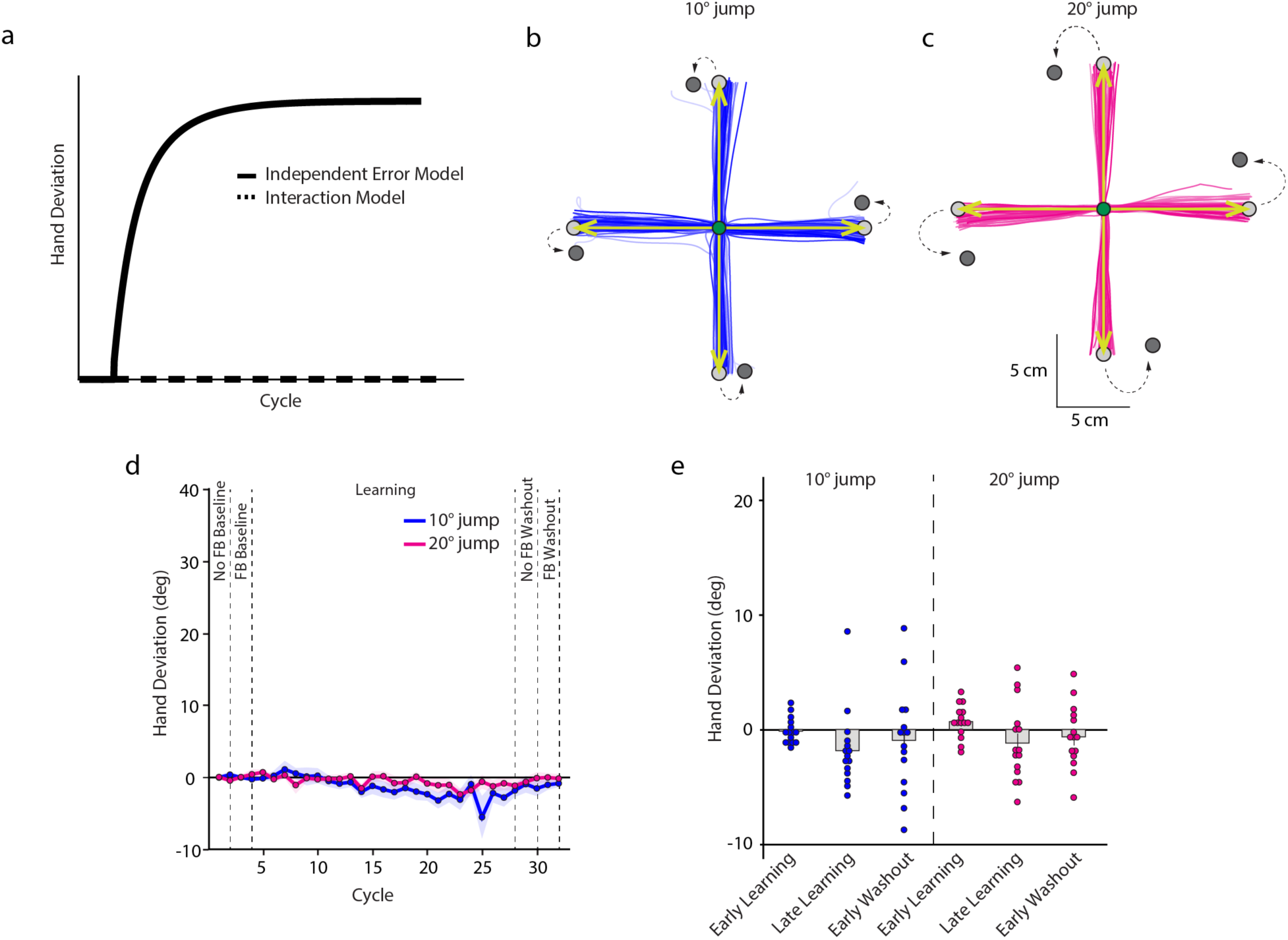
(a) Predictions of the Independent Error model and Interaction model for Experiment 2. Since there is no SPE in target displacement, the interaction model would predict no learning as shown by the dotted line. The Independent Error model, represented by the solid line, suggests normal learning. Hand trajectories of example subjects in the **(b)** 10° and **(c)** 20° target displacement groups during the learning block. Earlier trials are in lighter shades. The first 10 cm of the movement are shown. The green arrows indicate the clamped cursor trajectory. **(d)** Group-averaged hand angle deviation across cycles of different blocks. Shaded regions denote SEM. **(e)** Mean hand deviation during the early and late learning stages, as well as the early washout phase. Dots represent individual subjects.

Remarkably, our experimental data revealed little change in reach direction. As can be seen in Figs 2b and 2c, the hand was almost always directed towards the original target, even though it had been extinguished and a new one was displayed at a location 10° (Fig 2b) or 20° (Fig 2c) clockwise from it. The absence of a change in reach direction was consistent in the group-averaged data as well (Fig 2d). During early learning, hand deviation (Fig 2e) remained close to zero for the 10° (−0.154±0.29°, CI = [-0.777, 0.468]) as well as the 20° (0.687±0.378°, CI = [-0.123, 1.498]) groups. This continued to be the case at the end of learning as well (Fig 2e; 10°: - 1.843±0.887°, CI = [-3.747, 0.06], 20°: −1.182±0.863°, CI = [-3.033, 0.67]). Furthermore, after-effects on the early no-feedback washout trials were absent in both groups (Fig 2e; 10°: - 0.951±1.183°, CI = [-3.488, 1.585]; 20°: −0.648±0.708°, CI = [-2.167, 0.871]), which continued to be the case in the second washout block with visual feedback (10°: −1.051±0.702°, CI = [-2.556, 0.453]; 20°: −0.033±0.612°, CI = [-1.345, 1.278]).

Thus, when instructed to ignore performance failures, no adaptive changes in reach behavior occurred; this would *not* be the case if these errors were driving implicit tuning of reach plans. This result effectively ruled out the Independent Error framework as an explanation of the findings from Experiment 1. In other words, Experiment 2 showed that TPEs *cannot* independently induce implicit learning; rather an SPE is required for implicit learning to be set in motion.

### Experiment 3

The results of Experiment 2 help us disambiguate the Independent Error and the Interaction models and appear to support the latter. However, they may not directly substantiate the Interaction model. This is because no SPE is present in this experiment, and therefore any automatic modulation of SPE-driven learning in the presence of a TPE cannot be ascertained. In order to decisively test whether SPE-driven learning is influenced by TPEs, we performed a third experiment in which two groups of subjects reached under 30° counterclockwise error clamp conditions. Notably, for one of the groups (“Clamp+Jump”), the reach target was also jumped 30° counterclockwise on each learning trial, which caused the clamped cursor to always strike it at the end of the stipulated movement distance. Since the cursor always hit the target, no TPE was present. In the other group (“Clamp”), the target remained stationary and thus the (clamped) cursor always failed to strike it, resulting in a TPE. This arrangement then allowed us to probe if and how the SPE-mediated learning induced by the error clamp would change in the presence / absence of a TPE. Note that in this case, the Interaction model predicts differences in the way the hand angle would change for the Clamp and Clamp+Jump groups (Fig 3a). More specifically, the model suggests attenuation of learning in the Clamp+Jump group which does not experience TPEs, but a large change in hand angle for the Clamp group which does experience them.

**Fig 3.**
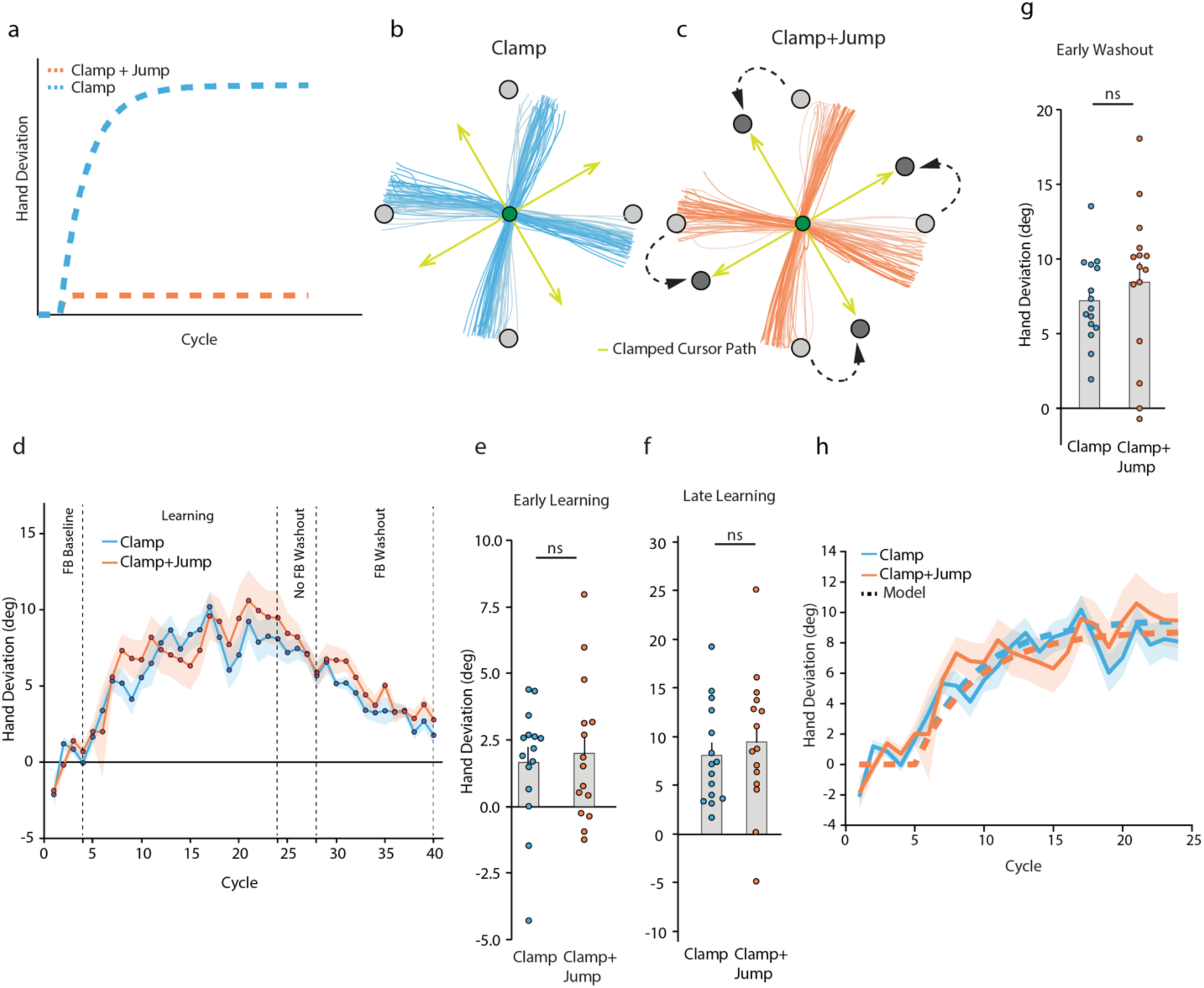
**(a)** Predictions of the Interaction model for Experiment 3. *This model predicts a robust change in hand angle for the Clamp group, but attenuated learning for the Clamp*+*Jump group*. Hand trajectories of example subjects of the **(b)** Clamp and **(c)** Clamp+*Jump groups during the learning block*. *Earlier trials are in lighter shades*. *The first 10 cm of the movement are shown*. *The cursor path is indicated by green arrows*. **(d)** Group averaged hand deviation (*relative to the original target direction*) *across cycles of the experimental blocks*. *Shaded regions denote SEM*. Hand deviation during the **(e)** early learning, **(f)** late learning and (**g**) early no-*feedback washout phases*. *Dots represent individual participants*. **(h)** Interaction model fits to the data of experiment 3. *Solid lines represent the experimentally observed hand deviation* (*shaded region denotes SEM*)*, while the dashed lines show the model fits*.

Remarkably, our experimental data revealed no group differences. A clear shift in the hand angle over the learning phase was evident in both groups (Figs 3b and 3c), which was confirmed in the group data as well (Fig 3d). There were no group differences in hand deviation during early learning (Fig 3e; Clamp: 1.653±0.580°, Clamp+Jump: 1.999±0.685°, t(27.261) = - 0.385, p = 0.703, Cohen’s d = −0.141). By the end of learning, both groups showed robust changes in hand angle (Fig 3f; Clamp: 8.087±1.323°, t(14) = −4.839, p < 0.001, Cohen’s d = −1.249; Clamp+Jump: 9.464±1.836°, t(14) = −4.081, p < 0.001, Cohen’s d = −1.054), but again, group differences were not significant at this time point (t(28) = −0.608, p = 0.548, Cohen’s d = - 0.222). As was the case in Experiment 1, there was no association between the change in hand angle from late baseline to early learning and the change over the rest of the learning phase (Fig S2b; Clamp: R^2^ = 0.127, p = 0.192; Clamp+Jump: R^2^ = 0.053, p = 0.409).

During the early no-feedback washout trials, large after-effects were evident in both groups. Notably, hand deviation in the Clamp group on the early after-effect trials was not different from late learning (Fig 3g; mean = 7.185±0.752°, t(14) = −1.114, p = 0.284, Cohen’s d = −0.288), which was also the case for the Clamp+Jump group (mean = 8.434±1.336°, t(14) = −1.069, p = 0.303, Cohen’s d = −0.276). Critically, a direct comparison of this early after-effect magnitude between the two groups indicated no significant differences (t (28) = −0.815, p = 0.422, Cohen’s d = −0.298). Furthermore, after-effects persisted even when cursor feedback was restored (Clamp: 6.559 ± 0.533°, CI = [5.415, 7.703]; Clamp+Jump: 6.744 ± 1.017° CI = [4.563, 8.926]), with no significant between-group differences (t(21.16) = −0.161, p = 0.873, Cohen’s d = −0.059). Remarkably, hand deviation did not reach baseline levels even on the last washout cycle for either group (Clamp: 1.762±0.626°, CI = [0.420, 3.103]; Clamp+Jump: 2.779±0.655°, CI = [1.373, 4.184]).

In sum, the experimental data revealed that despite experiencing a TPE, the overall performance of the Clamp group was not different from that of the Clamp+Jump group, which had not encountered any TPE.

#### Interaction model fit

We then fit the Interaction model (equation 4) to the data from Experiment 3 to probe for any potential groups difference in model parameters (Fig 3h). First, we fit the model to the mean data of the Clamp group while holding G_A_ and G_B_ at 1. We obtained a good fit (R^2^ = 0.8242, RMSE = 1.3519), and model parameters were estimated as A_SPE_ = 0.9199 and B_SPE_*SPE = 0.7181. We then used these parameter values and fit the model to the data from the Clamp+Jump group to estimate G_A_ and G_B_. Note that if the estimated values of G_A_ and G_B_ from this latter fit differ significantly from 1, it would imply that the TPE experienced by the Clamp+Jump group has some influence on the SPE-mediated learning induced by the error clamp. However, we obtained G_A_ and G_B_ estimates as 0.9986 (CI = [0.9667,1.0292]) and 1.1019 (CI = [0.6118,1.6640]) respectively (confidence intervals derived from fits to 10000 bootstrap samples of data from this group). This suggested that the TPE did not produce any modulatory effect.

### Experiment 4

Data from our first three experiments and our modeling efforts suggested that TPEs neither trigger implicit learning (Experiments 1 and 2) nor does their presence automatically modulate implicit learning mediated by SPEs (Experiment 3). How then do TPEs contribute? A final possibility that we considered was that TPEs induce strategic re-aiming to rapidly reduce the imposed errors. In Experiment 4, two groups of subjects made reaching movements under conditions in which cursor feedback was always veridical with the hand, thereby eliminating any SPE. However, one group experienced a 10° counterclockwise target jump, while the other experienced a 30° clockwise jump with respect to the original target, thus creating TPEs early on. Subjects in both groups were instructed to reach to the on-screen target, but at the end of learning and just before the after-effects block, the 30° jump group received the additional instruction that the target would stop jumping and that they should bring their hand to the original target location. This instruction was not given to the 10° jump group. We predicted that subjects in both groups would show increased RT and fleeting after-effects, classic signatures of a re-aiming strategy (23).

Fig 4a shows the hand trajectories of example subjects from the 10° and 30° jump groups respectively, while Fig 4b shows the group-averaged change in hand angle over the course of the experiment. It is evident that on the learning trials, the hand was directed towards the displaced target within a few trials, but did not go substantially beyond it. For both groups, there was a clear change in hand angle (relative to late baseline) early on itself (Fig 4c; 10°: t(9) = 3.76, p = 0.0045, Cohen’s d = 1.189; 30°: t(14) = 5.352, p<.001, Cohen’s d = 1.382). There was also a robust increase in RT during this early phase (Fig 4d; 10°: 96.17±24 ms; 30°: 229.27±44.55 ms), a change that was statistically significant (10°: t(9) = 4.015, p = 0.003, Cohen’s d = −1.27; 30°: t(14) = 5.146, p < 0.001, Cohen’s d = 1.329). By the end of learning, the mean change in hand angle relative to the late baseline stage was 9.014±0.653° and 28.883±1.008°, both of which were remarkably close to the magnitude of the jumps that the two groups experienced. Furthermore, in sharp contrast to Experiments 1 and 3, the early changes in hand angle were negatively correlated with changes over the rest of the learning block (Fig S2c; 10°: R^2^ = 0.587, p = 0.0097; 30°: R^2^ = 0.8, p < 0.001). In other words, if the initial change in hand angle was large enough to offset the error, no further learning occurred.

**Fig 4.**
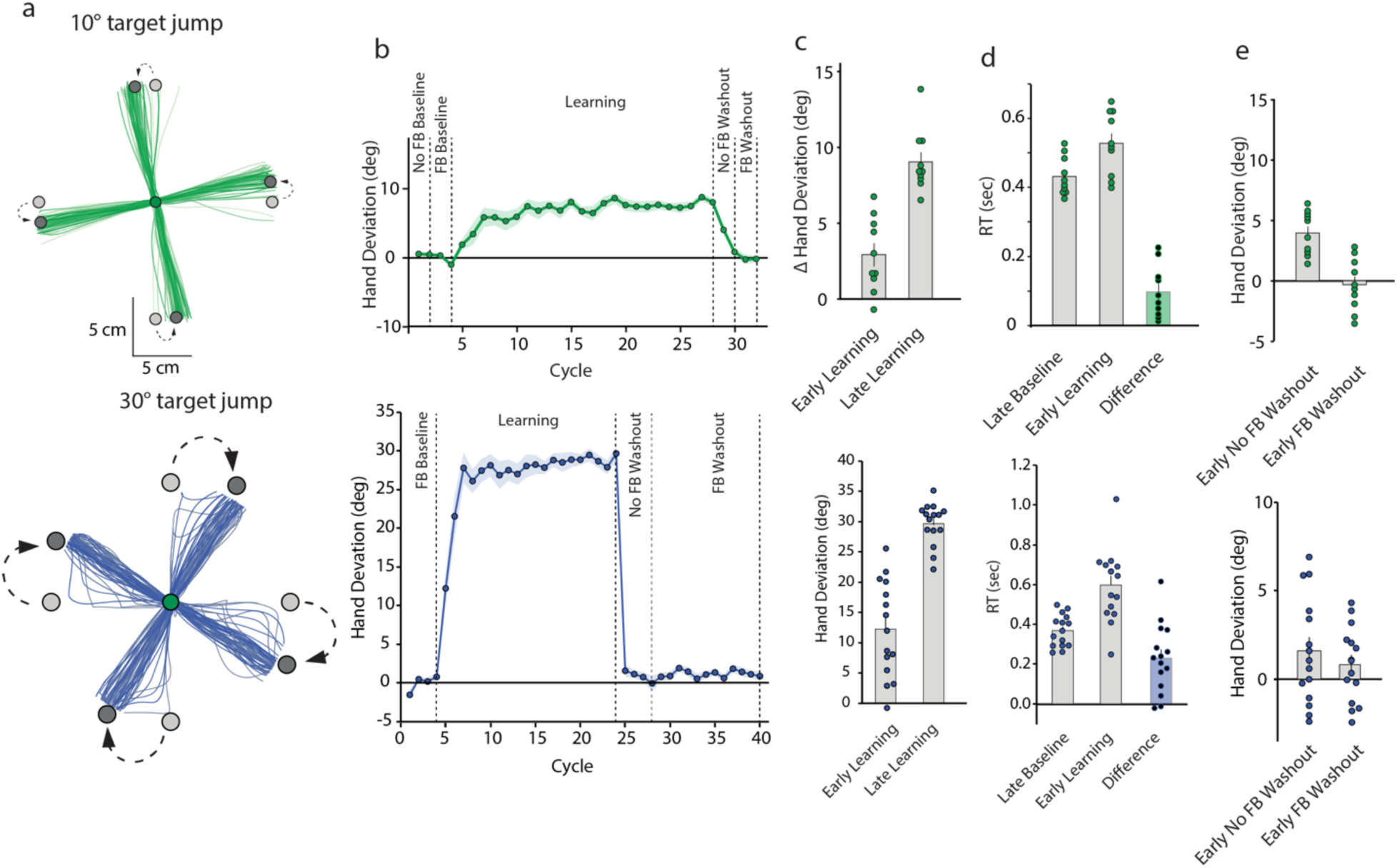
**(a)** Hand trajectories of example subjects during the learning block. Earlier trials are in lighter shades. The first 10 cm of the movement are shown**. (b)** Group averaged hand deviation across cycles of different experimental blocks. Shaded regions are SEM**. (c)** Change in hand angle during early and late learning relative to the late baseline stage. Dots are individual participants. **(d)** Group-averaged RT during the late baseline and early learning phases**. (e)** Group-averaged hand deviation on the early no-feedback and feedback washout trials. In all panels, the upper row represents 10° target jump group while the lower row represents 30° target jump group.

An interesting dichotomy emerged in the after-effects of the two groups on the early no-feedback washout trials (Fig 4e). For the 10° jump group, the hand remained somewhat deviated early on (mean = 3.95±0.566°, CI = [2.667, 5.227]), but for the 30° jump group, there was an immediate return to baseline (mean = 1.595±0.777°, CI = [-0.072, 3.262]). By the end of the first washout block however, hand deviation of even the 10° group reached near zero (mean = 0.757±0.526°, CI = [-0.434, 1.948]) and the 30° group continued to maintain similar levels (mean = −0.028±1.002, CI = [-2.178, 2.122]). There was no further change in hand angle over the remaining washout trials and hand deviation at the end of the feedback washout trials hovered around zero for both groups (10°: mean = −0.245±0.452°, CI = [-1.266, 0.777]; 30°: 0.900±0.654, CI = [-0.503, 2.303]). This suggested that after-effects of both groups were highly labile, but the 30° group showed a very rapid return to baseline since they were able to disengage from the strategy upon instruction prior to the start of the washout block. Additional analyses involving a direct comparison of this decay in these two TPE-only groups to that of the Hit and Miss groups of Experiment 1 confirmed the fleeting nature of the after-effects (Fig S3). This result, complemented with the robust increase in RT on the early learning trials, suggested that subjects accounted for the jump-induced TPEs via time-consuming re-aiming strategies that were rapidly disengaged when they were no longer relevant.

## Discussion

Error is believed to be the currency that drives sensorimotor adaptation to novel task conditions. We investigated the influence of different error signals on adaptive changes in motor output; we asked how limb-related SPEs and task-related performance failures (TPEs) contribute to learning. Our findings are at odds with the notion that performance failures bring about implicit recalibration of action plans (20, 21); rather, we observe that implicit learning is engaged only by SPEs. We also do not find evidence that SPE-driven learning is obligatorily modulated by the occurrence of TPEs (22). Rather, TPEs appear to set in motion distinct, verbally-sensitive strategic action selection to rapidly offset the error. The combination of TPE-driven strategies and SPE-driven implicit recalibration then likely determines how much and how rapidly our future action plans are tuned based on errors that we experience.

A hallmark of implicit recalibration is its imperviousness to verbal instruction (3, 6). That is, people *will* demonstrate implicit learning even when they are explicitly asked to ignore the error. However, in contrast to insensitivity, we found remarkable enslaving of TPE-induced responses to verbal directives. Consider the subjects who were instructed to *respond* to the TPE by aiming to a new target location – the Miss group of Experiment 1 and the two jump groups of Experiment 4. All these participants demonstrated an increase in RT and a rapid change in hand direction on the early learning trials, both of which are signatures of deliberative strategy use to offset the error. Furthermore, during washout, the participants of Experiment 4, who experienced only jump-induced TPEs, showed transient (10° jump) or negligible (30° jump) after-effects, indicating almost immediate disengagement from the learned behavior, another robust feature of strategy use. Even more remarkable was the performance of subjects in Experiment 2, who were instructed to *ignore* the TPE. In complete compliance with the instruction, these subjects showed a remarkable lack of change in reach direction and no after-effects. Taken together, these findings refute the notion that TPEs, at least those induced via target shifts, independently induce implicit recalibration, and strongly advocate that responses to TPEs involve deliberate, strategic selection of actions that bring about a change in movement direction. This ability to either deploy or disengage strategies “on call”, despite being a time-consuming process, provides a remarkably powerful means to adjust motor behavior to task demands. While the specific nature of these strategies remains to be elucidated, potential candidates include volitional, goal-directed control (23) or mental rotation of the original movement plan (24, 25).

Besides compensating for performance failures as shown here, recent work has revealed another major advantage of strategy use during learning. Notably, employing deliberative re-aiming could be the gateway to long-term storage of updates made to actions plans and expression of the acquired memory as savings. In particular, strategies that lead to rewarding task outcomes could be reinforced and recalled later, resulting in faster learning upon re-exposure to the original learning conditions (26, 27). In contrast, if strategy formation is prohibited by limiting movement planning time, by preventing exposure to performance errors, or by forcing only implicit learning, savings is blocked (28, 29). This suggests that unsuccessful task outcomes and associated strategy use are essential for forming long-term motor memories.

While our results argue that TPEs stimulate volitional strategy use, a pertinent question is why don’t TPEs induce implicit recalibration? One possibility is that for recalibration to occur, the sensorimotor system must attribute the error to some internal source, such as a limb-related execution failure (30, 31). It is plausible that TPEs, particularly those induced via target shifts, are instead seen as resulting from an external cause (30). The concomitant failure to accomplish the task goal could then be seen as arising from an action selection error (rather than an execution error), which then only requires the selection of a different action on the next trial (i.e., re-aiming) rather than updates to our internal models of the body and the world. In case of consistently occurring TPEs, action selection could be enhanced by learning task structure (extracting as much information about the environment as possible) instead of engaging the slower implicit system (23). Once such learning has occurred, actions could be guided by representations of the outcomes they produce and what these outcomes are worth, which, in fact, is precisely what is advocated in model-based reinforcement learning. It is also known that such algorithms can be adjusted rapidly to account for outcome revaluation as well as changes in the evironment and action goals (23, 32, 33). The quick, instruction-driven disengagement and return to baseline behavior on the after-effect trials of Experiment 4 is well-aligned with this idea, and suggests that “learning” from TPEs could proceed in this fashion.

Our current findings indicate that implicit learning is set in motion only when an SPE is present. In the presence of a concurrent TPE that subjects are expected to respond to, this implicit learning likely rides on top of the strategic adjustments to reach direction that the TPE sets in motion. This is precisely what we believe is the case with the Miss subjects of Experiment 1. As we noted earlier, our data from this group suggest that their early response involves explicit re-aiming to the shifted target location. This re-aiming then creates an SPE as people expect the cursor to follow the hand being aimed to the new target, while it remains clamped in the direction of the original target. The occurrence of this SPE in turn sets in motion implicit recalibration, which dominates to bring about further changes in hand angle and produce after effects (also see point 3 in Supplementary Information). Note that this SPE is just like a “classic” SPE that occurs as people move in an instructed direction while feedback about that motion is shown in another. However, our novel task design enables us to create it “on the fly” by requiring the Miss subjects to (first) respond to the TPE. Importantly, in this scheme, when strategy use is not employed – either due to verbal instruction or the absence of TPEs – net updates to action plans are determined entirely by the implicit process. This likely explains the attenuated learning in the Hit group of Experiment 1 as well as the Clamp and Clamp+Jump subjects in Experiment 3, and is perhaps also the basis for the reduced learning seen in other studies in which performance errors are eliminated (18–20).

Relatedly, in Experiment 2, although subjects experience a TPE like the Miss group of Experiment 1, they are asked to ignore it and reach the orignal target location. These subjects do not re-aim and no SPE is created since the hand and the clamped cursor are both directed towards the original target. Since no SPE occurs, no adaptive response occurs even though TPEs are present. These findings are reminiscent of classic work showing that in the presence of prediction errors, humans compulsively update their action plan in an implicit manner even if such learning bears a cost on performance errors (6). In line with this result, stroke patients with lesions to right inferior frontal and dorsolateral prefrontal cortices, who demonstrate clear performance failures when exposed to a perturbation, have no problem implicitly updating their reach direction in response to SPEs (34). Tellingly, recent work (35) suggests that the prediction error dominance may be so strong that task outcomes may have negligible influence, at least in canonical learning paradigms. Our results also strengthen this view.

Is it possible that implicit recalibration engaged by SPEs is modulated by the occurrence of TPEs as a default? The results of Experiment 3 do not favor this possibility. We found, consistent with past work (29), that neither the rate nor the amount of implicit learning was influenced by the presence (Clamp group) or absence (Clamp+Jump group) of TPEs. It may be that instead of the consistent and binary present / absent, or successful / unsuccessful conditions employed in the current study, trial-to-trial changes in SPE and TPE magnitudes might cause an influence of one on the other (22). However, some very recent results have not found any modulatory effects using more or less the same experimental paradigm (36). It also appears unlikely that modulation of SPE-mediated learning requires that TPEs are induced using conditions in which the target remains stationary. Indeed, the target location did not change for subjects in the Clamp group, yet their performance was not different from the group that experienced no TPEs (Clamp+Jump group). As such, evidence for direct modulation of SPE-driven learning through binary TPEs remains scant.

This is not to say however that the presence of binary TPEs cannot “influence” the net learning (change in hand angle). As our results themselves point out, such an influence arises only when subjects are expected to respond to TPEs. This is specifically reflected in the fact that although both groups experienced TPEs, the hand deviation of the Miss group of Experiment 1 (which was expected to respond to the target shift and aim to the new target) was nearly 3 times larger than that of the Clamp subjects of Experiment 3 (who were instructed to ignore the jump and move their hand to the original target location). We believe that the requirement to respond triggered strategic adjustment of aiming direction in the Miss group initially (followed by SPE-mediated implicit adjustments, as noted above) resulting in a much larger net change than the Clamp group. Barring such an instruction, the TPE carries no influence, and net learning is driven completely by the SPE, which is presumably what transpired in the Clamp group. Our results thus point to a dichotomy in mechanisms that the two error sources stimulate.

The distinction between processes that limb-related SPEs and goal-related TPEs stimulate suggests that these mechanisms are likely neurally separable as well. While prediction-error-based implicit learning depends on the cerebellum (2, 3) and parietal cortex (1, 14, 37), performance failures activate reward-sensitive cortico-striatal pathways (30, 38). Failure to obtain reward could trigger re-aiming via processes dependent on M1 and premotor cortex (39, 40). Changes in these regions following learning (41, 42) may thus partially reflect changes in action plans driven by such processes. Future work could probe the communication between these two systems, which, neuroanatomically, could be sustained by reciprocal connections between the basal ganglia and the cerebellum (43).

## Materials and Methods

### Participants

A total of 115 healthy, right-handed adults (85 males, 30 females, age range: 18-40) participated in the study. None of the participants reported any neurological, orthopedic or cognitive impairments. All subjects gave written informed consent and were monetarily compensated for their time. The project was approved by the Institute Ethics Committee of the Indian Institute of Technology Gandhinagar.

### Apparatus

The experimental setup consisted of a virtual reality system wherein participants sat facing a digitizing tablet and used a stylus to make hand movements on it (Fig S1a). A high-definition display was mounted horizontally above the tablet and was used to show circular start positions and targets for the reach, as well as a feedback cursor that would typically indicate the hand (stylus) location on the tablet. Participants looked into a mirror which was placed between the display and the tablet, and which reflected the display screen. The mirror also functioned to block direct vision of the arm. This arrangement enabled us to dissociate motion of the feedback cursor from that of the hand. For instance, cursor feedback could be veridical with the hand, “clamped” in certain directions independent of hand movement direction, or eliminated altogether.

### Task Procedure and Experimental Design

The general task involved making center-out reaching movements from a fixed start circle to a target. To initiate a trial, participants first moved their hand (cursor) into the start circle. After 500 ms, the reach target was displayed along with a beep, which indicated to subjects that they should begin moving. No constraint was placed on reaction time or movement time. Targets were presented at a distance of 10 cm and could appear at one of four locations arranged radially around a virtual circle in 90-degree increments (0°, 90°, 180°, 270°). The order of target appearance was pseudo-random; a target appeared in one of the four locations only once over four consecutive trials. This order was maintained across all participants. Cursor feedback, whenever provided, was shown continuously (throughout the movement) for a distance of 10 cm at which point the cursor “froze” and stayed in place even though the hand could continue moving.

#### Experiments 1 and 2

In Experiments 1 and 2, subjects performed 3 continuous blocks of trials: baseline, learning and washout. The baseline and washout blocks comprised of two sub-blocks each. In the first “no-feedback” baseline sub-block (20 trials), the cursor was not shown during the reach, while in the second “feedback” sub-block (20 trials), the cursor was shown veridical with the hand. Following baseline, subjects experienced 240 learning trials, which were followed by the two washout sub-blocks that were identical to baseline (20 no-feedback, 20 veridical feedback trials). Each subject thus performed 320 trials in all. During the baseline and washout blocks, the originally displayed target (at one of the 0°, 90°, 180°, 270° locations) remained “stationary”, i.e. its location did not change during the trial. During the learning block however, the target was displaced, or “jumped”, by 10° in the counterclockwise direction on every trial. This was achieved by extinguishing the originally presented target as the hand reached mid-way to it, and displaying a new target at the 10°, 100°, 190° or 280° locations. Importantly, on these trials, the motion of the feedback cursor was always “clamped” in the direction of the original target. In other words, the cursor always followed a direct, straight path to the original target regardless of the direction of hand motion.

##### Experiment 1

In Experiment 1, a start circle of 0.9 cm diameter and targets of 0.98 cm diameter were used. Participants were randomly divided into two groups, “Miss” or “Hit” (n=15 each), which differed in terms of the learning trials experienced. For the Miss group, the original and the new target were of the same size, but for the Hit group, the diameter of the new target was increased from the original 0.98 cm to 4.6 cm (Fig S1b). Since cursor motion was always clamped towards the original target, the target jump resulted in the cursor missing the new target for the Miss group, thereby resulting in a task performance failure (TPE). However, for the Hit participants, the increase in target size ensured that the cursor would still hit the new target, thus preventing the task error. This hit was obviously not in the center of the new target.

Subjects were explained the task conditions prior to start of the experiment. They were instructed to reach towards the target on the screen. For baseline and washout trials, this obviously meant the original, fixed target. However, for learning trials, since the original target was extinguished and a new one displayed in an easily-predictable location, subjects could start aiming in the direction of the new target. No attempt was made to either dissuade or encourage this behavior. In addition, subjects were also informed and made to understand that on the jump (learning) trials, cursor motion would be fixed and would not depend on the direction of their hand movements. They were also explicitly told to ignore this cursor feedback. A reminder to this effect was provided at the halfway point of the learning block.

##### Experiment 2

The overall task design for Experiment 2 remained identical to Experiment 1. We recruited two groups of subjects (n=15 each), who experienced target jumps of different amplitudes (10° or 20°) while the cursor remained clamped towards the original target. This led to conditions similar to the Miss group of Experiment 1 where the cursor did not strike the new, shifted target thus creating TPEs. Critically however, both groups of subjects were now also explicitly instructed to ignore the change in target location and reach towards the original target location. As in Experiment 1, subjects were reminded of this halfway into the learning block.

#### Experiment 3

In Experiment 3, subjects performed 400 center-out reaching movements from a start circle (1.2 cm diameter) to four different targets (1.5 cm diameter) in three continuous blocks: baseline (40 trials with veridical cursor feedback), learning (200 trials), and washout (40 no-feedback and 120 veridical feedback trials). In the learning block, the cursor was visible, but was rotated and clamped at 30° counterclockwise relative the target direction. That is, the cursor always followed a fixed, rotated path relative to the target independent of the direction of the underlying hand motion.

Participants of Experiment 3 were randomly divided into two groups, “Clamp” and “Clamp+Jump” (n=15 each) that differed in terms of the learning trials experienced. While the 30° cursor clamp was enforced for both groups during learning, the reach target was also jumped for the “Clamp+Jump” participants to a location that was 30° counterclockwise to the original target location. This was not the case for the “Clamp” group, for whom the original target remained “stationary” and visible on the screen throughout. Given this arrangement, the Clamp subjects always experienced both an SPE and a TPE since the rotated, clamped cursor never hit the target. In contrast the Clamp+Jump participants experienced an SPE due to the clamp, but no TPE since the cursor always landed on the shifted target at the end of the stipulated movement distance. The Clamp subjects were instructed to ignore the cursor and reach to the on-screen target, while the “Clamp+Jump” subjects were additionally asked to ignore the shift in target location and continue reaching to the original target location.

#### Experiment 4

In our fourth experiment, subjects once again performed three blocks of centre-out reaching trials: baseline, learning and washout. Subjects were randomly divided in two groups. One of the groups (n=10) performed 40 baseline trials (20 no-feedback, 20 feedback), 240 learning trials and 40 washout trials (20 feedback, 20 no-feedback) from a start circle (0.9 cm diameter) to four different targets (0.98 cm diameter). The second group performed 40 feedback baseline trials, 200 learning trials and 160 washout trials (40 no-feedback, 120 feedback) from a start circle of 1.2 cm diameter to targets of 1.5 cm diameter. For both groups, cursor feedback, when shown, was always veridical with the hand. However, subjects in the first group experienced a 10° counterclockwise target jump on the learning trials, while subjects in the second groups experienced a 30° clockwise target jump. Given that the cursor was always veridical with the hand, no SPE occurred. However, the target shift would induce a consistent TPE during the learning block in both groups.

Participants in both groups were instructed to reach to the target on the screen. This meant the originally displayed target on the baseline and washout trials, and the new, shifted target on the learning trials. Importantly, at the end of the learning block and prior to start of the washout block, participants in the 30° jump group were given the additional instruction that the target would stop jumping and that they should reach to the original target location. Such an instruction was not given to the 10° jump group.

### Mathematical modelling

We used different variants of the state-space model framework of motor adaptation to better understand our behavioral findings, particularly those of Experiment 1, and make predictions about subsequent experiments (20). The model equations are typically of the following form:

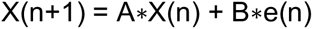

where *e* represents the error on the n^th^ trial, X represents the motor output, and A and B represent a retention factor and error sensitivity respectively.

#### Independent error model

First, we considered a framework in which both sensory prediction errors (SPEs) and performance errors (TPEs) drive independent implicit processes. The net motor output reflects the sum of these SPE- and TPE-based processes. The model equations are:

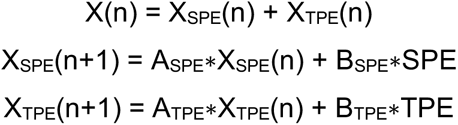

In Experiment 1, in the Hit condition, the TPE-based update does not occur (since TPE = 0) and learning can be presumed to be driven only by the SPE-based process. However, for the Miss case, both processes get updated since the SPE and TPE are both present, and the net output is the sum of these two processes.

We first fit only the SPE-sensitive process of the independent error model to the cycle-wise data of the Hit group using the *fmincon* function of Matlab. Additionally, B_SPE_*SPE was estimated as a single parameter since the SPE remained fixed due to the clamp. A_SPE_ was the other parameter estimated from the fit. Data from the baseline, learning and no-feedback washout blocks were included, with the B_SPE_*SPE term being set to zero for the no-feedback washout cycles. The feedback washout data were not used since the SPE was no longer constant in this condition (clamp was removed) and could change as subjects changed their hand direction. The estimated values of A_SPE_ and B_SPE_*SPE derived from model fits to the Hit participants’ data were then used while fitting the model that included the TPE-based process to the data of the Miss participants of Experiment 1. Thus, only A_TPE_ and B_TPE_*TPE terms were estimated from these latter fits. For both fits, A_SPE_ and A_TPE_ were constrained between 0 and 1, while B_SPE_*SPE and B_TPE_*TPE were constrained between 0 and 10°. The 10° value corresponds to the maximum “error” (angle between new target direction and clamped cursor direction) on any given trial.

#### Interaction model

In our second “Interaction” model, TPEs cannot by themselves induce implicit learning, but can only modulate implicit learning induced by SPEs. The equations governing the trial-by-trial updates to motor output can be given by:

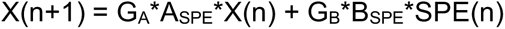

Here, X represents the internal state, SPE signifies the sensory prediction error, A and B are the retention factors and error sensitivity respectively, and G_A_ and G_B_ represent parameters that modulate them. When fitting to the data from Experiment 1, for the Miss group, G_A_ and G_B_ can be set as 1, while they can be estimated from model fits for the Hit group (alternatively, G_A_ and G_B_ can also be set as 1 for the Hit group and estimated for the Miss group). Further, since the SPE remains constant in the error clamp, B*SPE is estimated as a single term.

For Experiment 1, we first fit the model to the data of the Miss group and estimated the values of A_SPE_ and B_SPE_*SPE (G_A_ and G_B_ were set to 1). We then used these estimates when fitting the model to the data of the Hit group and estimated the values of G_A_ and G_B_. The same procedure was followed in Experiment 3. The model was first fit to the data of the Clamp group (with G_A_ and G_B_ set to 1) and the values of A_SPE_ and B_SPE_*SPE were estimated. These values were then used when fitting the model to the Clamp+Jump group to estimate G_A_ and G_B_.

### Empirical Data Analysis and Statistics

Hand position data (X-Y coordinates) were filtered using a low-pass Butterworth filter with 10-Hz cutoff frequency. Velocity values were obtained by differentiating the position data. The primary dependent variable was the deviation in hand direction relative to the original target direction. This was computed as the angle between the line connecting the start position to the original target, and the line connecting the start position to the hand position at peak movement velocity.

Trials in which participants did not initiate a movement or lifted the stylus off the tablet mid-trial leading to loss of data were marked as “bad trials” and excluded from the analysis. Additionally, trials in which hand deviation was more than 85° were also removed. Collectively, across the 115 participants, 1.41% trials were excluded. We then calculated baseline directional biases, defined as the mean hand deviation across all baseline trials. These biases were subtracted from the trial-wise hand deviation data.

Learning was quantified using the cycle-by-cycle values of the baseline subtracted hand deviation over the learning block (1 cycle = 10 trials). For each subject, early learning was defined as the mean deviation over the first 10 learning trials while late learning was characterized by the mean deviation over the last 10 learning trials. Reaction time (RT) during early learning was assessed in terms of a change from late baseline levels. This was done by subtracting the mean RT of the last 10 baseline trials from the mean RT of the first 10 learning trials. Early after-effect magnitude was quantified as the mean deviation over the first 10 trials of the no-feedback washout block. To assess whether they were sustained, average after-effect magnitude on the first 10 trials of the feedback washout block was also calculated.

Group differences in hand deviation during early and late learning as well as the different after-effect stages were compared using Welch’s t-tests after checking for normality with Shapiro-Wilk tests. Paired t-tests were used for assessing changes in hand deviation across different time points within a group. Significance levels were set at 0.05. Cohen’s *d* was used for estimating the effect size of the differences. We also used the 95% confidence interval to probe for significant deviation in hand angle during early and late learning as well as the different after-effect stages.

## Author Contributions

AO, AK, AS and PKM designed the experiments. AO and AS collected the data, all the authors analyzed the data and interpreted the results. All authors also wrote and approved the final manuscript.

## Acknowledgments

This work was partially supported by grants from the Department of Science and Technology, Government of India to PKM. We also acknowledge support from IIT Gandhinagar. We thank Gaurav Panthi for helpful discussions.

## SUPPLEMENTARY INFORMATION

### 1. Experimental Setup

**Figure S1.**
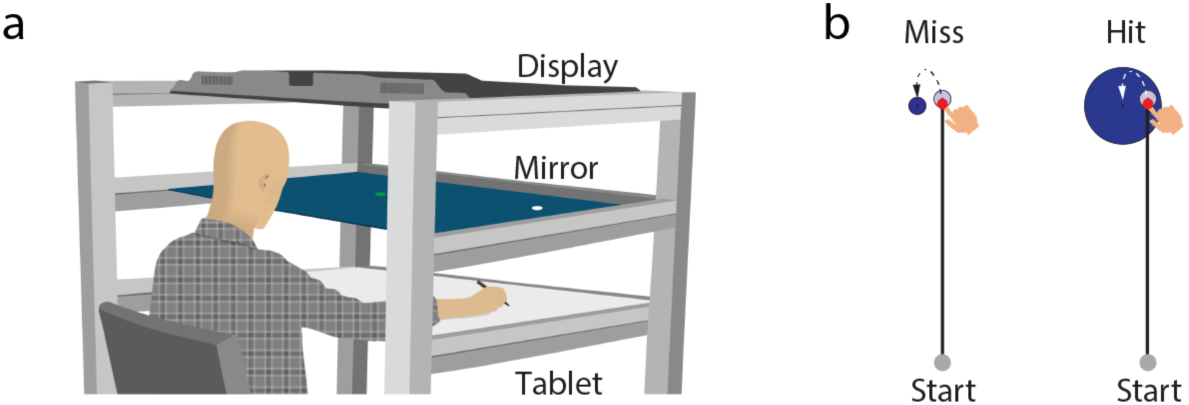
**(a)** Subjects performed reaching movements on a digitizing tablet using a handheld stylus while looking into a mirror placed between the tablet and a horizontally mounted display screen. Movements were restricted to the horizontal plane. Start positions, targets, and a feedback cursor were displayed on the screen, and were reflected in the mirror**. (b)** On learning trials, the target was displaced, or **“**jumped**”** 10° counter**-**clockwise, while cursor motion was clamped in the direction of the original target. For the Miss group, this resulted in a performance failure since the cursor failed to strike the new target. For the Hit group, target size was increased along with the jump, which caused the clamped cursor to always hit the new target despite the shift in target location.

### 2. Relationship between early and late changes in hand angle

The Miss group of experiment 1 experienced performance failures. During the early learning stage, these subjects demonstrate larger RTs and increased hand deviation (relative to baseline). As our experiment 4 and other work (1, 2) shows, such changes reflect the deployment of strategic processes that cause the hand to aim away from the original target. Importantly however, this early strategy use has no bearing on how their hand angle changes over the remainder of the learning block.

**Figure S2.**
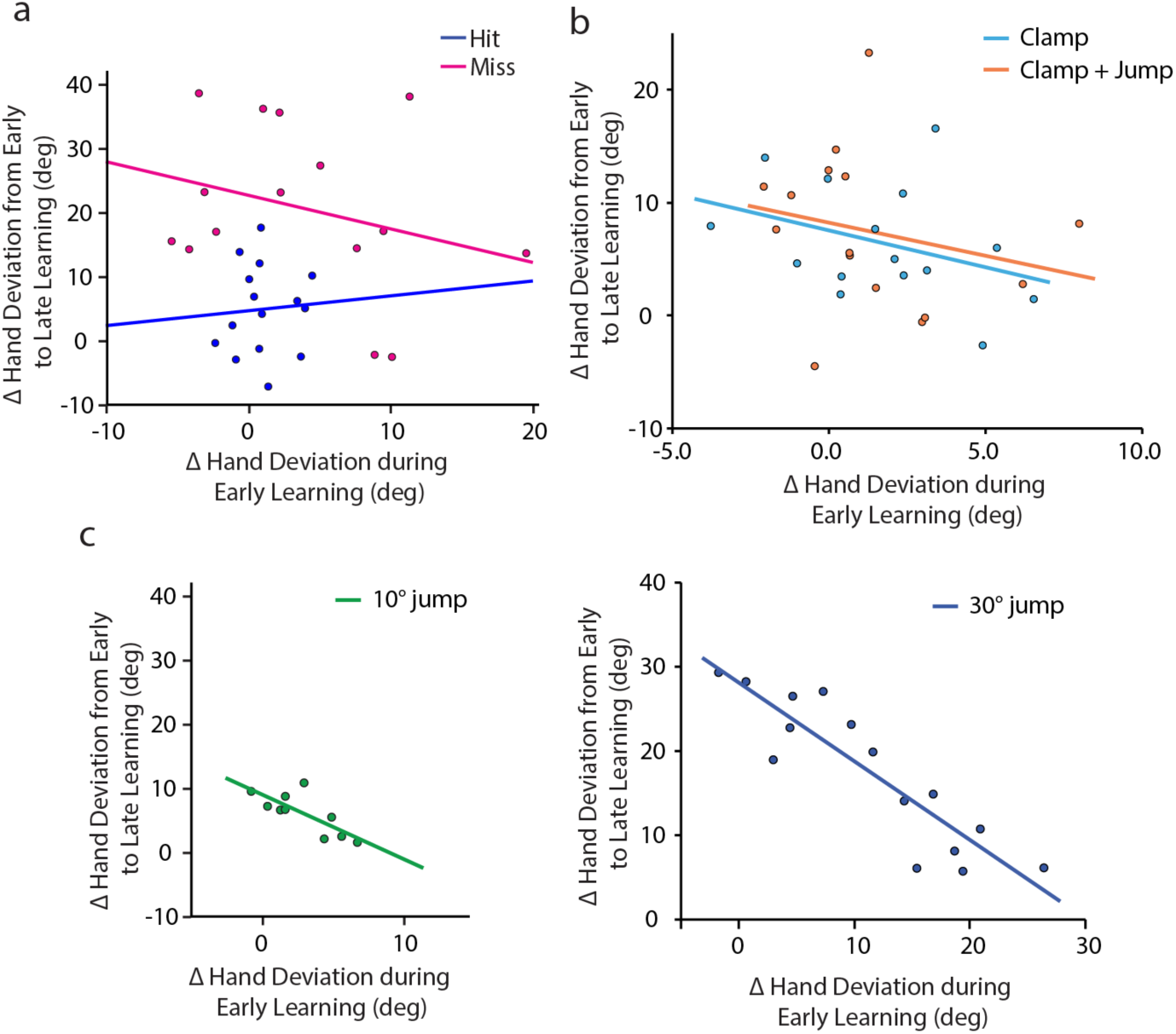
Relationship between the change in hand deviation from baseline to early learning, and the change in hand deviation over the rest of the learning block (***a***) *for hit and miss groups of experiment 1*, (***b***) *the Clamp and Clamp*+*Jump groups of experiment 3, and* (***c***) *the 10°* (*left*) *and 30° jump* (*right*) *groups of experiment 4*.

As Figure S2a (pink) shows, there is no relationship between the change in hand angle during early learning and the change thereafter until the end of learning. Our data suggest that the latter changes are driven by unconscious, implicit mechanisms that are engaged as subjects intend to aim to the new target location while cursor motion is clamped in the direction of the original target. Put together, our data for the Miss group suggest that the intent to aim to the new target emerges via explicit re-aiming with implicit mechanisms taking over thereafter (see elaboration in the Discussion section of the main manuscript).

The relationship between the early and later changes in hand deviation is also not seen in the Hit group (Figure S2a, blue). While on the face of it, this seems similar to the Miss group, the underlying reason is likely to be different. The Hit participants do not experience performance failures, and do not show any substantial change in hand angle or RT during the early learning stage. Thus, unlike the Miss group, these participants do not employ deliberative mechanisms to change their aim towards the new target early on. The changes that emerge over the course of the learning block then are likely driven entirely by an implicit process that is engaged as they gradually start aiming to the center of the new target while the cursor remains clamped in the original target direction. Thus, the absence of a relationship between early and late changes in the Hit group is because a single process that emerges slowly is engaged.

Likewise, the subjects of experiment 3 demonstrated no relationship between early deviation in hand angle and changes over the rest of the learning block (Figure S2b). The reason here is plausibly similar to that of the Hit participants of experiment 1. It is likely that subjects in both the Clamp and the Clamp+Jump groups employ a single, slower, implicit process to learn, as has been suggested in other reports that have engaged similar groups of subjects as well (3). Since early changes in hand angle are very small, and pretty much all learning occurs later, no relationship emerges between the early and subsequent changes in hand deviation. In sharp contrast, a strong negative relationship between the early change in hand angle and subsequent changes is seen for the groups in Experiment 4 (Figure S2c). Note that these groups do not experience an SPE, and changes in hand angle are driven only by a TPE induced by the target jump. The observed relationship therefore indicates that if the early process “siphons off” a large chunk of the error, little additional change in hand angle occurs. We suggest, based on the consistent increase in RT and hand angle during early learning as well as the labile after-effects, that this process is a conscious, re-aiming strategy. Since the error is compensated via this mechanism and a prediction error is not present, an additional (implicit) learning mechanism is not engaged.

### 3. Direct comparison of after-effects in experiments 1 and 4

The Hit and Miss groups of experiment 1 showed sustained post-learning after-effects. Subjects of experiment 4, who experienced only TPEs, showed deviations in hand angle immediately after the learning block had ended, but this effect was not sustained particularly for the group that experienced the 30° jump. However, a direct comparison between these groups is precluded since the asymptotic level of learning is different between them. To overcome this issue, we used the following approach. We considered only the hand angle data of the last learning cycle and the four washout cycles (two no-feedback, two feedback washout cycles). We then scaled the data of individual Hit and Miss participants of experiment 1, and the subjects who experienced the 10° jump in experiment 4 to match the mean hand angle of the 30° jump group on the last learning cycle. Thus, at the end of this step, the mean hand angle at the end of learning for all four groups was identical. We then compared the (scaled) after-effects across the four groups (Figure S3).

**Figure S3:**
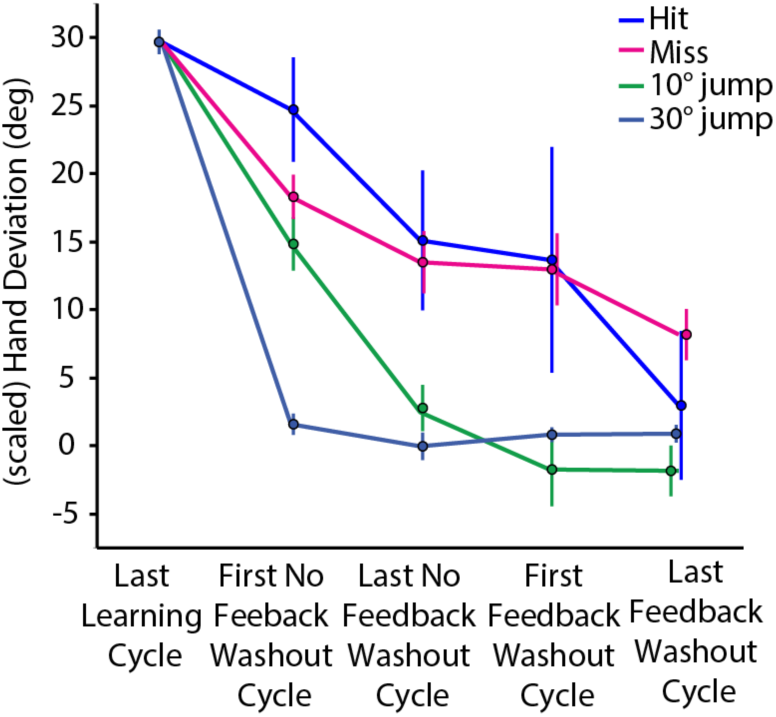
Direct comparison of after-effects in experiment 1 and experiment 4. Scaled average hand deviation of the Hit and Miss groups of experiment 1 and the 10° *and 30*° *jump groups of experiment 4. Data on the last learning cycle, the two no-feedback washout cycles and the two feedback washout cycles are shown. Error bars represent SEM. The (scaled) hand deviation on washout trials decayed more rapidly towards zero for the subjects of experiment 4*.

We found a clear group difference in the hand deviation on the first no-feedback washout cycle (F(3,51) = 17.671, p < 0.001, η^2^_p_ = 0.510). A Dunnett’s post-hoc test showed that relative to the 30° jump group, the three other groups, i.e., 10° jump (p=0.002), Miss (p<0.0001), and Hit (p<0.0001) had much larger had deviation. Group differences were also prevalent during the later no-feedback washout trials (F(3,51) = 5.918, p = 0.001, η^2^_p_ = 0.258), but Dunnett’s test now indicated a significantly greater hand angle for only the Hit (p = 0.0027) and Miss (p = 0.0080) groups, but not the 10° jump group (p = 0.9196) relative to the 30° jump group. This suggested that subjects in the 10° jump group had also returned to near baseline levels of hand deviation by this time. The group differences began to diminish as washout trials progressed; there was no major difference between the groups on the early feedback washout trials (F(3,51) = 2.592, p = 0.063, η^2^ = 0.132), and group differences were ngliglible at the end of washout (F(3,51) = 1.613, p = 0.198, η^2^ = 0.087).

As we have argued in the main manuscript, the subjects in experiment 4 (10° and 30° jump) compensated for the imposed TPE by employing a deliberative re-aiming strategy. As such, this strategy (of aiming away from the original target) would be irrelevant on washout trials. The highly transient nature of after-effects suggests that strategy use was rapidly “abandoned” by these subjects when it was no longer relevant. This was particularly the case for the 30° jump group that in fact was explicitly instructed before the washout trials that the target would stop jumping and they should move to the original target location. The 10° jump group also shows a rapid decline, with their hand deviation returning to baseline levels by the second washout cycle (this group was not given any instruction before washout). However, the Hit and Miss groups return to baseline levels much more slowly since their learning is dominated by SPE-driven implicit mechanisms.

## References

1. A. Kumar, G. Panthi, R. Divakar, P. K. Mutha, Mechanistic determinants of effector-independent motor memory encoding. Proc Natl Acad Sci U S A 117, 17338–17347 (2020).

2. T. A. Martin, J. G. Keating, H. P. Goodkin, A. J. Bastian, W. T. Thach, Throwing while looking through prisms. I. Focal olivocerebellar lesions impair adaptation. Brain 119 ( Pt 4, 1183–1198 (1996).

3. J. R. Morehead, J. A. Taylor, D. E. Parvin, R. B. Ivry, Characteristics of Implicit Sensorimotor Adaptation Revealed by Task-irrelevant Clamped Feedback. J Cogn Neurosci 29, 1061– 1074 (2017).

4. R. L. Sainburg, C. Ghez, D. Kalakanis, Intersegmental dynamics are controlled by sequential anticipatory, error correction, and postural mechanisms. J Neurophysiol 81, 1045–1056 (1999).

5. R. Shadmehr, F. A. Mussa-Ivaldi, Adaptive representation of dynamics during learning of a motor task. J Neurosci 14, 3208–3224 (1994).

6. P. Mazzoni, J. W. Krakauer, An implicit plan overrides an explicit strategy during visuomotor adaptation. Journal of Neuroscience 26, 3642–3645 (2006).

7. Y. W. Tseng, J. Diedrichsen, J. W. Krakauer, R. Shadmehr, A. J. Bastian, Sensory prediction errors drive cerebellum-dependent adaptation of reaching. J Neurophysiol 98, 54–62 (2007).

8. M. Synofzik, A. Lindner, P. Thier, The cerebellum updates predictions about the visual consequences of one’s behavior. Curr Biol 18, 814–818 (2008).

9. A. Clark, Whatever next? Predictive brains, situated agents, and the future of cognitive science. Behavioral and Brain Sciences 36, 181–204 (2013).

10. S. D. McDougle, K. M. Bond, J. A. Taylor, Explicit and Implicit Processes Constitute the Fast and Slow Processes of Sensorimotor Learning. J Neurosci 35, 9568–9579 (2015).

11. H. E. Kim, J. R. Morehead, D. E. Parvin, R. Moazzezi, R. B. Ivry, Invariant errors reveal limitations in motor correction rather than constraints on error sensitivity. Commun Biol 1 (2018).

12. K. Wei, K. Körding, Relevance of error: What drives motor adaptation? J Neurophysiol 101, 655–664 (2009).

13. K. M. Bond, J. A. Taylor, Flexible explicit but rigid implicit learning in a visuomotor adaptation task. J Neurophysiol 113, 3836–3849 (2015).

14. P. K. Mutha, R. L. Sainburg, K. Y. Haaland, Left parietal regions are critical for adaptive visuomotor control. J Neurosci 31, 6972–6981 (2011).

15. V. Della-Maggiore, N. Malfait, D. J. Ostry, T. Paus, Stimulation of the posterior parietal cortex interferes with arm trajectory adjustments during the learning of new dynamics. J Neurosci 24, 9971–9976 (2004).

16. M. Maschke, C. M. Gomez, T. J. Ebner, J. Konczak, Hereditary Cerebellar Ataxia Progressively Impairs Force Adaptation during Goal-Directed Arm Movements. J Neurophysiol 91, 230–238 (2004).

17. J. A. Taylor, J. W. Krakauer, R. B. Ivry, Explicit and implicit contributions to learning in a sensorimotor adaptation task. Journal of Neuroscience 34, 3023–3032 (2014).

18. L. A. Leow, W. Marinovic, A. de Rugy, T. J. Carroll, Task errors contribute to implicit aftereffects in sensorimotor adaptation. European Journal of Neuroscience 48, 3397–3409 (2018).

19. K. Van der Kooij, L. O. Wijdenes, T. Rigterink, K. E. Overvliet, J. B. J. Smeets, Reward abundance interferes with error-based learning in a visuomotor adaptation task. PLoS One 13 (2018).

20. H. E. Kim, D. E. Parvin, R. B. Ivry, The influence of task outcome on implicit motor learning. Elife 8 (2019).

21. S. T. Albert, et al., Competition between parallel sensorimotor learning systems. Elife 11 (2022).

22. J. S. Tsay, A. M. Haith, R. B. Ivry, H. E. Kim, Interactions between sensory prediction error and task error during implicit motor learning. PLoS Comput Biol 18 (2022).

23. D. P. Sadaphal, A. Kumar, P. K. Mutha, Sensorimotor Learning in Response to Errors in Task Performance. eNeuro 9 (2022).

24. J. Fernandez-Ruiz, W. Wong, I. T. Armstrong, J. R. Flanagan, Relation between reaction time and reach errors during visuomotor adaptation. Behavioural Brain Research 219, 8–14 (2011).

25. S. D. McDougle, J. A. Taylor, Dissociable cognitive strategies for sensorimotor learning. Nat Commun 10, 40 (2019).

26. D. M. Huberdeau, A. M. Haith, J. W. Krakauer, Formation of a long-term memory for visuomotor adaptation following only a few trials of practice. J Neurophysiol 114, 969–977 (2015).

27. J. R. Morehead, S. E. Qasim, M. J. Crossley, R. Ivry, Savings upon re-aiming in visuomotor adaptation. Journal of Neuroscience 35, 14386–14396 (2015).

28. A. M. Haith, D. M. Huberdeau, J. W. Krakauer, The influence of movement preparation time on the expression of visuomotor learning and savings. Journal of Neuroscience 35, 5109– 5117 (2015).

29. L. A. Leow, W. Marinovic, A. de Rugy, T. J. Carroll, Task errors drive memories that improve sensorimotor adaptation. Journal of Neuroscience 40, 3075–3088 (2020).

30. J. Diedrichsen, Y. Hashambhoy, T. Rane, R. Shadmehr, Neural correlates of reach errors. J Neurosci 25, 9919–9931 (2005).

31. J. S. Tsay, H. Kim, A. M. Haith, R. B. Ivry, Understanding implicit sensorimotor adaptation as a process of proprioceptive re-alignment. Elife 11 (2022).

32. N. D. Daw, Y. Niv, P. Dayan, Uncertainty-based competition between prefrontal and dorsolateral striatal systems for behavioral control. Nat Neurosci 8, 1704–1711 (2005).

33. B. B. Doll, D. A. Simon, N. D. Daw, The ubiquity of model-based reinforcement learning. Curr Opin Neurobiol 22, 1075–1081 (2012).

34. P. K. Mutha, R. L. Sainburg, K. Y. Haaland, Critical neural substrates for correcting unexpected trajectory errors and learning from them. Brain 134, 3647–61 (2011).

35. J. Wang, S. Bao, G. D. Tays, Lack of generalization between explicit and implicit visuomotor learning. PLoS One 14, e0224099 (2019).

36. N. Al-Fawakhiri, A. Ma, J. A. Taylor, O. A. Kim, On the money and right on target: How robust are reward and task success effects on implicit motor adaptation? bioRxiv, 2023.02.01.526533 (2023).

37. P. K. Mutha, L. H. Stapp, R. L. Sainburg, K. Y. Haaland, Motor Adaptation Deficits in Ideomotor Apraxia. Journal of the International Neuropsychological Society 23, 139–149 (2017).

38. M. Inoue, M. Uchimura, S. Kitazawa, Error Signals in Motor Cortices Drive Adaptation in Reaching. Neuron 90, 1114–1126 (2016).

39. A. P. Georgopoulos, J. T. Lurito, M. Petrides, A. B. Schwartz, J. T. Massey, Mental rotation of the neuronal population vector. Science (1979) 243, 234–236 (1989).

40. S. M. Kosslyn, G. J. DiGirolamo, W. L. Thompson, N. M. Alpert, Mental rotation of objects versus hands: Neural mechanisms revealed by positron emission tomography. Psychophysiology 35, 151–161 (1998).

41. R. Paz, T. Boraud, C. Natan, H. Bergman, E. Vaadia, Preparatory activity in motor cortex reflects learning of local visuomotor skills. Nat Neurosci 6, 882–890 (2003).

42. M. G. Perich, J. A. Gallego, L. E. Miller, A Neural Population Mechanism for Rapid Learning. Neuron 100, 964–976.e7 (2018).

43. A. C. Bostan, R. P. Dum, P. L. Strick, The basal ganglia communicate with the cerebellum. Proc Natl Acad Sci U S A 107, 8452–8456 (2010).

## Supplementary References

1. J. Fernandez-Ruiz, W. Wong, I. T. Armstrong, J. R. Flanagan, Relation between reaction time and reach errors during visuomotor adaptation. Behavioural Brain Research 219, 8–14 (2011).

2. S. D. McDougle, J. A. Taylor, Dissociable cognitive strategies for sensorimotor learning. Nat Commun 10, 40 (2019).

3. L. A. Leow, W. Marinovic, A. de Rugy, T. J. Carroll, Task errors drive memories that improve sensorimotor adaptation. Journal of Neuroscience 40, 3075–3088 (2020).

